# Hitting more birds with one stone: CD70 as an actionable immunotherapeutic target in recurrent glioblastoma

**DOI:** 10.1101/2021.06.02.446670

**Authors:** M Seyfrid, W Maich, MV Shaikh, N Tatari, D Upreti, D Piyasena, M Subapanditha, N Savage, D McKenna, L Kuhlmann, A Khoo, SK Salim, B Bassey-Archibong, W Gwynne, C Chokshi, K Brown, N Murtaza, D Bakhshinyan, P Vora, C Venugopal, J Moffat, SK Singh

**Affiliations:** Departments of Surgery, Faculty of Health Sciences, McMaster University, 1200 Main Street West, Hamilton, Ontario, L8N 3Z5, Canada; Biochemistry and Biomedical Sciences, Faculty of Health Sciences, McMaster University, 1200 Main Street West, Hamilton, Ontario, L8N 3Z5, Canada; Princess Margaret’s Hospital, Toronto, Ontario, M5S 3E1, Canada; Donnelly Centre, Department of Molecular Genetics, University of Toronto, Ontario, Canada

**Keywords:** Glioblastoma, CD70, Immunotherapy, CAR-T

## Abstract

**Purpose:** Glioblastoma (GBM) patients suffer from a dismal prognosis, with standard of care therapy inevitably leading to therapy-resistant recurrent tumors. The presence of brain tumor initiating cells (BTICs) drives the extensive heterogeneity seen in GBM, prompting the need for novel therapies specifically targeting this subset of tumor-driving cells. Here we identify CD70 as a potential therapeutic target for recurrent GBM BTICs.

**Experimental Design:** In the current study, we identified the relevance and functional influence of CD70 on primary and recurrent GBM cells, and further define its function using established stem cell assays. We utilize CD70 knockdown studies, subsequent RNAseq pathway analysis, and *in vivo* xenotransplantation to validate CD70’s role in GBM. Next, we developed and tested an anti-CD70 CAR-T therapy, which we validated *in vitro* and *in vivo* using our established preclinical model of human GBM. Lastly, we explored the importance of CD70 in the tumor immune microenvironment (TIME) by assessing the presence of its receptor, CD27, in immune infiltrates derived from freshly resected GBM tumor samples.

**Results:** CD70 expression is elevated in recurrent GBM and CD70 knockdown reduces tumorigenicity in vitro and in vivo. CD70 CAR-T therapy significantly improves prognosis *in vivo*. We also found CD27 to be present on the cell surface of multiple relevant GBM TIME cell populations.

**Conclusion:** CD70 plays a key role in recurrent GBM cell aggressiveness and maintenance. Immunotherapeutic targeting of CD70 significantly improves survival in animal models and the CD70/CD27 axis may be a viable poly-therapeutic avenue to co-target both GBM and its TIME.

## INTRODUCTION

Glioblastoma (GBM) is the most common malignant brain tumor in adults accounting for approximately 14.6% of all brain tumors (Ostrom et al. 2019). Despite an aggressive standard of care (SoC) including maximal surgical resection and chemoradiotherapy, GBM patients have a median survival time of less than 15 months, and a five-year survival rate of less than 6.8% (Stupp et al. 2005; Stupp et al. 2017; Ronning et al. 2012). GBM often recurs 7-9 months after resection of the primary tumor, at which point the tumor is often non-resectable, and poorly responsive to chemo- and/or radiotherapy, leaving patients with therapeutic options limited to clinical trial enrollment (van Linde, M. E., 2017).

In the past three decades, survival rates across several cancers have improved significantly, due in part to major advances in technology allowing for early detection, as well as significant leaps in targeted and novel therapeutic strategies (Siegel, Miller, and Jemal 2018). However, despite these advances, little to no improvement has been made in prognosis for GBM patients, who continue to suffer from dismal outcomes.

Therapeutic failure, in part, is due to extensive intratumoral heterogeneity at the cellular, genetic, and functional levels (Bergmann et al. 2020; Soeda et al. 2015; Xiong, Yang and Li 2020). This heterogeneity may be explained by a distinct subset of cells coined brain tumor initiating cells (BTICs) (Soeda et al. 2015), which possess stem cell-like traits such as self-renewal, therapy evasion, and multi-lineage differentiation (Cusulin et al. 2015). It is believed that this subpopulation of BTICs, after undergoing selective pressures from primary GBM SoC therapy, become chemo- and radio-therapy resistant, and seed formation of the therapy-resistant recurrent tumors (Liu et al. 2006; Bao et al. 2006). Expression of GBM BTIC markers such as CD133, CD15, and CD44 are generally associated with worse clinical outcome (Zeppernick et al. 2008). Thus, novel therapeutic interventions to target not only the tumor bulk, but the treatment resistant BTIC population that seeds recurrence is necessary.

Immunotherapy holds great promise in cancer treatment, and recent studies in gliomas provide encouraging results (Vora et al. 2020; Morgan et al. 2012; Jin et al. 2018). Amongst various immunotherapeutic approaches are adoptive T cell therapies, including chimeric antigen receptor (CAR) T-therapy. CAR-Ts are T-cells expressing a recombinant cell-surface receptor that directs these cells to specific tumor associated antigens (TAAs). Upon binding to the TAA, T-cells undergo MHC-independent activation and induce apoptosis of the target cell (Priceman, Forman, and Brown 2015; Zhang et al. 2017). However, to develop safe and effective CAR-T cells, novel tumor-specific antigens with a sufficient therapeutic window are required.

Genomic and proteomic data from a multi-omic target development pipeline revealed CD70 as a suitable therapeutic target in recurrent GBM (rGBM). Our data shows that CD70 is more highly expressed in rGBM samples compared to primary GBM (pGBM) samples. Moreover, CD70 is absent on normal human astrocytes and neural stem cells, as is supported by the literature (Grewal et al. 2008), thereby presenting a novel opportunity to target recurrent GBM. CD70 is a transmembrane glycoprotein and a member of the tumor necrosis factor (TNF) superfamily, and is the only known ligand for CD27. While CD70 is transiently expressed on activated T- B-cells, as well as mature dendritic cells, it is minimally expressed in most normal tissues (Jacobs et al. 2015, Adam P.J. et al. 2006). Similarly, CD27 is primarily only expressed on specific subsets of T- B-, and NK cells (Bowman et al. 1994; Hintzen et al. 1994; Tesselaar et al. 2003). The CD70/CD27 signalling axis leads to differentiation, proliferation, and T- and B-cell survival and proliferation (Denoeud and Moser 2011; Nolte et al. 2009; Borst, Hendriks, and Xiao 2005). Prolonged expression of CD70 has been shown to elicit lethal immunosuppression in mice (Tesselaar et al. 2003), and result in exhaustion of effector memory T-cells in B-cell non-Hodgkin lymphoma (Yang et al. 2014).

CD70 displays aberrant constitutive expression in a variety of cancers, include renal cell carcinoma, leukemia, non-small cell lung cancer, melanoma, GBM, and others (Diegmann et al. 2005; Lens et al. 1999; Jacobs et al. 2015; Pich et al. 2016; Held-Feindt and Mentlein 2002; Hishima et al. 2000; Pahl et al. 2015). In 2005, researchers showed that in B-cell lymphoma, CD70 and CD27 are mutually overexpressed, resulting in increased proliferation and survival of tumor cells via amplified signalling through the CD70/CD27 axis (Nilsson et al. 2005). In the context of GBM, CD70 has been shown to promote tumor progression and invasion (Ge et al. 2017). While in healthy individuals CD70 plays a role in eliciting an immune response, its role in the tumor microenvironment is far more multi-faceted. Within the GBM microenvironment, CD70 mediates immune escape (Wischhusen et al. 2002), and its overexpression leads to recruitment and activation of immunosuppressive T regulatory cells (Tregs) (Claus et al. 2012) and tumor associated macrophages (TAMs) (Ge et al. 2017). Together, these studies suggest that CD70 plays a major role in the recruitment and maintenance of the GBM immunosuppressive microenvironment, while promoting pro-tumorigenic processes.

The soluble form of CD27 (sCD27) is detected at high levels in the blood of cancer patients (Purdue et al. 2019; Kashima et al. 2019). Currently, there are multiple therapeutic strategies targeting CD70-expressing malignancies (Aftimos et al. 2017; Bristol-Myers Squibb 2013; Seagen Inc. 2018), however, the prevalence of the CD70/CD27 interaction provides a rationale for synergistic therapeutic opportunities targeting both tumor cells and the immune microenvironment.

To our knowledge, this is the first time CD70 has been identified as an immunotherapeutic target on BTICs from patient-derived GBM samples. In the presented work, we conduct a systematic study evaluating the efficacy of CD70 CAR-T cells in using our established patient-derived GBM mouse model, illustrating the potential of a CD70-directed CAR-T therapy to offer hope to GBM patients suffering from a dismal prognosis.

## Material and methods

### Dissociation and culture of primary GBM tissue

Human GBM samples (Table S1) were obtained from consenting patients, as approved by the Hamilton Health Sciences/McMaster Health Sciences Research Ethics Board. Brain tumor samples were dissociated in PBS (ThermoFisher, Cat#10010049) containing 0.2 Wunsch unit/mL Liberase Blendzyme 3 (Millipore Sigma, Cat#5401119001), and incubated in a shaker at 37°C for 15 min. The dissociated tissue was filtered through a 70μm cell strainer (Falcon, Cat#08-771-2) and collected by centrifugation (1500 rpm, 3 min). Red blood cells were lysed using ammonium chloride solution (STEMCELL Technologies, Cat#07850). GBM cells were resuspended in Neurocult complete (NCC) media, a chemically defined serum-free neural stem cell medium (STEMCELL Technologies, Cat#05751), supplemented with human recombinant epidermal growth factor (20ng/mL: STEMCELL Technologies, Cat#78006), basic fibroblast growth factor (20ng/mL; STEMCELL Technologies Cat#78006), heparin (2 mg/mL 0.2% Heparin Sodium Salt in PBS; STEMCELL technologies, Cat#07980), antibiotic-antimycotic (1X; Wisent, Cat# 450-115-EL), and plated on ultra-low attachment plates (Corning, Cat#431110) and cultured as neurospheres. GBM8 and GBM4 was a kind gift from Dr. Hiroaki Wakimoto (Massachusetts General Hospital, Boston, MA, USA), RN1, S2b2 and WK1 were gifts from Dr Andrew Boyd (QIMR Berghofer Medical Research Institute, Australia).

### Propagation of Brain tumor stem cells (BTSCs)

Neurospheres derived from minimally cultured (< 20 passages) human GBM samples were plated on polyornithine-laminin coated plates for adherent growth. Adherent cells were replated in low-binding plates and cultured as tumorspheres, which were maintained as spheres upon serial passaging *in vitro*. As shown before, compared to commercially available GBM cell lines, patient derived 3D cultures represent the variety of heterogeneous clones present within patient samples (Patrizii et al. 2018). These models recapitulate the key GBM morphological, architectural and expression features that are present in primary GBM. These cells retained their self-renewal potential and were capable of *in vivo* tumor formation.

### Glycocapture Proteomics

Briefly, cells were lysed in PBS:TFE (50:50) using pulse sonication and by incubating the lysates at 60°C for 2 hours (lysates were vortexed every 30 minutes). Protein concentration was determined using the BCA assay (Pierce). Cysteines were reduced with DTT (5mM final concentration) at 60°C for 30 minutes and alkylation was performed by adding iodoacetamide (25mM final concentration) to the cooled lysates and subsequent incubation at room temperature for 30 minutes. Trypsin was added at a 1:500 ration and protein digestion was performed overnight at 37°C. Tryptic peptides were desalted on C18 Macrospin columns (Nest Group), lyophilized and resuspended in coupling buffer (0.1M Sodium Acetate, 0.15M Sodium Chloride, pH 5.5). Glycan chains were oxidized using 10mM NaIO_4_ for 30 minutes in the dark and peptides were again desalted. Lyophilized peptides were resolubilized in coupling buffer and oxidized glycopeptides were captured on hydrazide magnetic beads (Chemicel, SiMAG Hydrazide) for 12h at room temperature. The coupling reaction was catalysed by adding aniline (50mM) and the reaction was allowed to continue for additionally 3 hours at room temperature.

Hydrazide beads containing the covalently coupled oxidized glycopeptides were thoroughly washed (2 x coupling buffer; 5x 1.5M NaCl; 5x HPLC H2O; 5x Methanol; 5x 80% Acetonitrile; 3x Water; 3x 100mM NH_4_OH, pH 8.0) to remove non-specific binders. N-glycopeptides were eluted off the hydrazide beads using 5U PNGase F in 100mM ammonium bicarbonate at 37°C overnight. The de-glycosylation reaction converts the asparagine residue, covalently linked to a glycan chain, to aspartic acid, the process carrying a signature mass shift of 0.98 Da.

Eluted (i.e. deamidated) glycopeptides were recovered and the hydrazide beads were additionally washed 2X with 80% acetonitrile solution. Glycopeptides were desalted using C18 stage tips, eluted using 80% acetonitrile, 0.1% F.A. and lyophilised. The purified glycopeptides were dissolved in 21μL 3% acetonitrile, 0.1% F.A. Peptide concentration was determined using a NanoDrop 2000 (Thermo) spectrophotometer.

### RNA sequencing and GSEA/Cytoscape analysis

Total RNA was extracted using the Norgen Total RNA isolation kit (Cat #48400) and quantified using a NanoDrop Spectrophotometer ND-1000. The RNA was sequenced using single-end 50 bp reads on the Illumina HiSeq platform (Illumina, San Diego CA, USA). Raw sequence data were exported to FASTQ format and were filtered based on quality scores (Quality cutoff of 20 for at least 90% of the bases in the sequence). Next the reads were mapped to the UCSC mRNA transcript human database based on the GRCh38/hg38 version using HISAT. The counts were obtained by using ht-seq count with the “intersection-strict” option. Counts were transformed with TMM transformation and then normalized with VOOM (package “limma” in R).

Differential gene expression profiles were generated by DESeq2 using the Galaxy online suite (https://usegalaxy.org/) and as imput of the Gene Set Enrichment Analysis (GSEA). Gene sets were randomized at 2000 permutations per analysis against Oncogenic (C6), Curated (C2) and Hallmark MSigDB collections of gene sets (https://www.gsea-msigdb.org/gsea/msigdb/index.jsp).

### Secondary sphere formation assay

Tumorspheres were dissociated using 5-10μL Liberase Blendzyme3 (0.2 Wunsch unit/mL) in 1mL PBS for 5 min at 37°C. Based on each cell line’s growth kinetics, cells were plated at 200-1000 cells per well in 200 μL of NCC media in a 96-well plate. Cultures were left undisturbed at 37°C, 5% CO2. After four days, the number of secondary spheres formed were counted.

### Cell proliferation assay

Single cells were plated in a 96-well plate at a density of 200-1000 cells/200 μL (based on each cell line’s growth kinetics) per well in quadruplicate and incubated for five days. 20 μL of Presto Blue (ThermoFisher, Cat#A13262), a fluorescent cell metabolism indicator, was added to each well approximately 4h prior to the readout time point. Fluorescence was measured using a FLUOstar Omega Fluorescence 556 Microplate reader (BMG LABTECH) at excitation and emission wavelengths of 535 nm and 600 nm respectively. Readings were analyzed using Omega analysis software.

### Receptor internalization and Antibody Drug Conjugate assay

For detection of internalization, 200,000 cells were being used for each condition, where both were incubated with antibody 30min on ice, rinsed twice and let incubated for 2h either at 37°C or at 4C, before being analyzed under flow cytometry.

GBM BTICs expressing CD70 on their cell surface were seeded and incubated for 30 minutes with different concentrations of he-lm-Fab’2 anti-CD70, followed by addition of 13nM of 2^◦^ADC *a*-HFab-NC-MMAF (conjugated with Monomethyl auristatin F) (Moradec, Cat# AH-121-AF) and proliferation was measured after 5 days (n=3 for BT241s, n=2 for HEK293s and n=1 for BT935s). The manufacturer’s protocol was followed directly, with the exception of using twice the initial amount of recommended antibody (40 nM).

### *In vivo* intracranial injections and H&E/immunostaining of xenograft tumors

Animal studies were performed according to guidelines under Animal Use Protocols of McMaster University Central Animal Facility. Intracranial injections in 6–8 week-old NSG mice were performed as previously described (Singh et al. 2004) using either BT241, GBM8 or GBM4 cells (100,000 cells/mice). Briefly, a burr hole is drilled at the point located 2 mm behind the coronal suture, and 3 mm to the right of the sagittal suture and GBM cells suspended in 10 μL PBS are intracranially injected with a Hamilton syringe (Hamilton, Cat#7635-01) into right frontal lobes of 6-8 week-old NSG mice. For CAR-T treatment, ConCAR-T or CD70CAR-T cells were injected intratumorally once a week for two weeks (for BT241, 1M first week then 0.5M; for GBM8 0.75M first week then 1M for GBM8). For tumor volume evaluation, animals were sacrificed when control mice reached endpoint. When mice reached endpoint, they were perfused with 10% formalin and collected brains were sliced at 2mm thickness using brain-slicing matrix for paraffin embedding and H&E staining. Images were captured using an Aperio Slide Scanner (Leica Biosystems) and analyzed using ImageScope v11.1.2.760 software (Aperio). For survival studies, all the mice were kept until they reached endpoint and number of days of survival were noted for *Kaplan Meyer* Analysis. CD3 stained slides were scanned and captured using an Aperio Slide Scanner and analyzed using ImageScope v11.1.2.760 software (Aperio). Tumor areas were generated using Aperio Membrane Algorithm.

### Generation of CAR Lentivirus

Human anti-CD70 (he_l and he_lm) scFv sequence were synthesized with a 5’ leader sequence and 3’ Myc tag by Genescript. The scFv was cloned into the lentiviral vector pCCL ΔNGFR (kindly provided by Dr. Bramson, McMaster University, Hamilton, ON, Canada) down-stream of the human EF1α promoter leaving ΔNGFR intact downstream of the minimal cytomegalovirus promoter. Empty pCCL ΔNGFR was used as a control vector. Replication-incompetent lentiviruses were produced by co-transfection of the CAR vectors and packaging vectors pMD2G and psPAX2 in HEK293FT cells using Lipofectamine 3000 (ThermoFisher, Cat#L3000075) as recommended by the manufacturer. Viral supernatants were harvested 24 and 48 hours after transfection and concentrated by ultracentrifugation at 15,000 RPM for 2h at 4°C. The viral pellet was resuspended in 1.0 mL of T cell media, aliquoted and stored at -80°C.

### Generation of CAR-T cells

Peripheral blood mononuclear cells (PBMCs) from consenting healthy blood donors were obtained using SepMate™ (STEMCELL technologies, Cat#85450). This research was approved by the McMaster Health Sciences Research Ethics Board. 1×10^5^ cells in XSFM media (Irvine Scientific, Cat#91141) were activated with anti-CD3/CD28 beads at a 1:1 ratio (Dynabeads, GIBCO, Cat#113.31D) in a 96-well round bottom plate with 100U/mL rhIL-2 (Peprotech, Cat#200-02). Twenty-four hours after activation, T cells were transduced with lentivirus at a MOI∼1. CAR-T cell cultures were expanded into fresh media (XSFM media supplemented with 100U/mL rhIL-2) as required for a period of 6-8 days prior to experimentation.

### Evaluation of cytokine release

NGFR+ sorted CAR-T cells (CD70CAR or ConCAR) were co-incubated with GBM cells at a 1:1 ratio for 24 hours. Supernatants were collected in duplicate for each condition and stored at 80°C for analysis of cytokines. Human TNF-α DuoSet ELISA kit (R & D Systems, Cat #: DY210-05) and IFN-γ DuoSet ELISA kit (R & D Systems, Cat #: DY285B-05) were used for quantification of the two cytokines by ELISA, according to manufacturer’s recommendation.

### Flow cytometric analysis and sorting

GBM cells and T cells in single cell suspensions were resuspended in PBS+2mM EDTA. GBM cells were stained with he_l or he_lm IgG (0.064-1000nM) or by IgG control AffiniPure Goat Anti-Human IgG, F(ab’)2 fragment specific, Jackson ImmunoResearch, Cat#109-005-006) or APC-conjugated anti-CD70 antibody (Miltenyi Biotech, REA 292) and incubated for 30 min on ice. CAR-T cells were stained with fluorescent tagged anti-CD3 (BD Biosciences, Cat#557851), anti-NGFR (Miltenyi Biotech, Cat#130-112-790) and anti-c-Myc (Miltenyi Biotech, Cat#130-116-653). Samples were run on a MoFlo XDP Cell Sorter (Beckman Coulter). Dead cells were excluded using the viability dye 7AAD (1:10; Beckman Coulter, A07704). Compensation was performed using mouse IgG CompBeads (BD Biosciences, Cat#552843).

### Cytotoxicity assays

Luciferase-expressing GBM cells at a concentration of 30,000 cells/well were plated in 96–well plates in triplicates. In order to establish the BLI baseline reading and to ensure equal distribution of target cells, D-firefly luciferin potassium salt (15 mg/mL) was added to the wells and measured with a luminometer (Omega). Subsequently, effector cells were added at 4:1, 3:1, 2:1, 1:1, and 0:1 effector-to-target (E: T) ratios and incubated at 37°C for 4-8 hours. BLI was then measured for 10 s with a luminometer as relative luminescence units (RLU). Cells were treated with 1% Nonidet P-40 (NP40, Thermofisher, Cat#98379) to measure maximal lysis. Target cells incubated without effector cells were used to measure spontaneous death RLU. The readings from triplicates were averaged and percent lysis was calculated with the following equation:

% *Specific lysis*=100(*spontaneous death RLU*− *test RLU*)(*spontaneous death RLU*−−*m aximal killing RLU*)%

Specific lysis=100(spontaneous death RLU− test RLU)(spontaneous death RLU—maxi mal killing RLU)

### Isolation and evaluation of immune cells from Brain tumor samples

EasySep human CD45 Depletion kit II (Stem Cell Technology, Cat#: 17898) was used to extract immune cells from freshly dissected patient tumors, according to the manufacturer’s protocol but with slight modifications. Briefly:

1. Prepare a single-cell suspension from tumor like before and resuspend the cells at **10^8cells/mL** in EasySep Buffer (Cat #: 20144) (If the tumor is too small, and total cells are less than 10^7 cells resuspend in 0.1 ml and adjust antibody and reagents (step 2 and 4 accordingly):
2. Add 12.5μL/mL of EasySep Human CD45 Depletion Cocktail II (Component#17898C)
3. Incubate 5min at RT
4. Add 20μL/mL of EasySep™ Dextran RapidSpheres 50101 (Component #50101)
5. Incubate 3min at RT
6. Top up volume (EasySep Buffer) to 2.5 ml and place tube into the EasySep magnet (Cat #: 18000) for 5 minutes
7. Pour off CD45-cells – cells remaining in tube are CD45+ population
8. For optimal recovery, perform 2 x 5min separation in the magnet

To identify individual tumor immune microenvironment immune cell populations, CD45+ cells were thawed and used immediately to run a panel of antibodies in order to identify individual immune cell populations. Antibodies used are as follows and were used according to manufacturer’s protocol: PE-Cy7 Mouse Anti-Human CD3 (Cat 563423; BD Pharmingen), PE Mouse Anti-Human CD4 (Cat555347; BD Pharmingen), APC Mouse Anti-Human CD8 (Cat555369; BD Pharmingen), PE-CF594 Mouse Anti-Human CD68 (Cat564944; BD Horizon), APC-H7 Mouse Anti-Human HLA-DR (Cat561358; BD Pharmingen), BV421 Mouse Anti-Human CD27 (Cat562514; BD Horizon).

### CAR-T Fratricide

Jurkat human T lymphocytes (Cedarlane Cat#: TIB-152) were expanded and grown in RPMI 1640 (Gibco Cat#:11875-093) with 10% FBS (Multicell Cat#:08105), 1% Penicillin-Streptomycin (Gibco Cat#:15140-122) and 10 mM HEPES (Gibco Cat#: 15630-080). Jurkat cells were then transduced with either an shGFP or shCD70 lentiviral construct and selected for by puromycin selection. shGFP Jurkat cells were sorted by flow cytometry to isolate a CD70^hi^ shGFP population, and untransduced Jurkat cells were sorted to isolate a CD70^hi^ population. shGFP and shCD70 Jurkat cells were then transduced with either Control CAR or CD70 CAR virus and allowed to expand. At 4 and 8 days after transduction, each population (shGFP ConCAR; shGFP CD70CAR; shCD70 ConCAR; shCD70 CD70CAR) was assessed by flow cytometry for the following markers: NGFR, CD70, CD69, viability.

### Statistical Analysis

Biological replicates from at least three patient samples were compiled for each experiment, unless otherwise specified in figure legends. Respective data represent mean ± SD, *n* values are listed in figure legends. Student’s t test analyses were performed using GraphPad Prism 5. p > 0.05 = n.s., p < 0.05 = *, p < 0.01 = **, p < 0.001 = ***, p < 0.0001 = ****.

## RESULTS

### CD70 expression is a unique marker of recurrent glioblastoma

Between the underrepresentation of rGBM samples in biobanks, due to the relatively low re-operation rate at GBM recurrence, and the variable presence of BTICs within bulk tumor samples, rGBM targets are often overlooked (Robin, Lee, and Kalkanis 2017). In this study, we leveraged an RNA sequencing platform using four in-house, low-passage BTIC-enriched cell lines derived from pGBM or rGBM patient samples, as previously described by our lab (Venugopal et al. 2012). Using the GBM TCGA repository (Bowman et al. 2017) we identified genes over-represented in BTIC-enriched populations, among which we found the TNF superfamily member CD70, which was present in pGBM BTICs but particularly upregulated in rGBM BTICs. (Fig1A, SupplFig1A). To further investigate the relevance of CD70 as a recurrent GBM marker, we used six primary/recurrent pairs from patient-matched GBM samples present in the TCGA database to evaluate CD70 expression. *In silico* analysis of CD70 mRNA expression revealed increased levels in rGBM samples compared to their matched primaries for the majority of the pairs available, however this trend did not reach significance (Fig1B). Additionally, these same matched pairs exhibited a Classical (TCGA-CL) to Mesenchymal (TCGA-MES) subtype transition from primary to recurrence, indicating a shift towards a more aggressive and therapy-resistant subtype with poorer prognosis (Wang et al. 2017, Sa et al. 2020). Given that mRNA expression does not necessarily translate directly to cell-surface protein expression, we interrogated cell-surface CD70 protein levels on two in-house matched primary/recurrent patient derived BTIC lines. We observed an increase in CD70 surface expression in both pairs by flow analysis (Fig1C) and a switch from the CL to MES subtype as seen in our bulk RNA sequencing samples (BT594/BT972, data not shown). We next screened a variety of unmatched primary and recurrent GBMs, as well as normal human cells lines (neural stem cells and astrocytes) for CD70 expression (Fig1D, SupplFig1B). We identified a therapeutic window with normal human brain cells, which minimally express CD70 on their cell surface, in accordance with the existing literature (Hintzen et al. 1994). We demonstrate a clear trend towards increased CD70 expression in rGBM compared to pGBM (Fig1C), which bulk tumor data corroborates (Rahman et al. 2018). Lastly, using our BT594/BT972 matched-pair, we performed N-Glycocapture Proteomics which ranked CD70 among the top upregulated cell surface markers in rGBM compared to pGBM (Fig1E). This data led us to further inquiries about the functional role that CD70 plays in GBM progression and maintenance.

**Figure 1:**
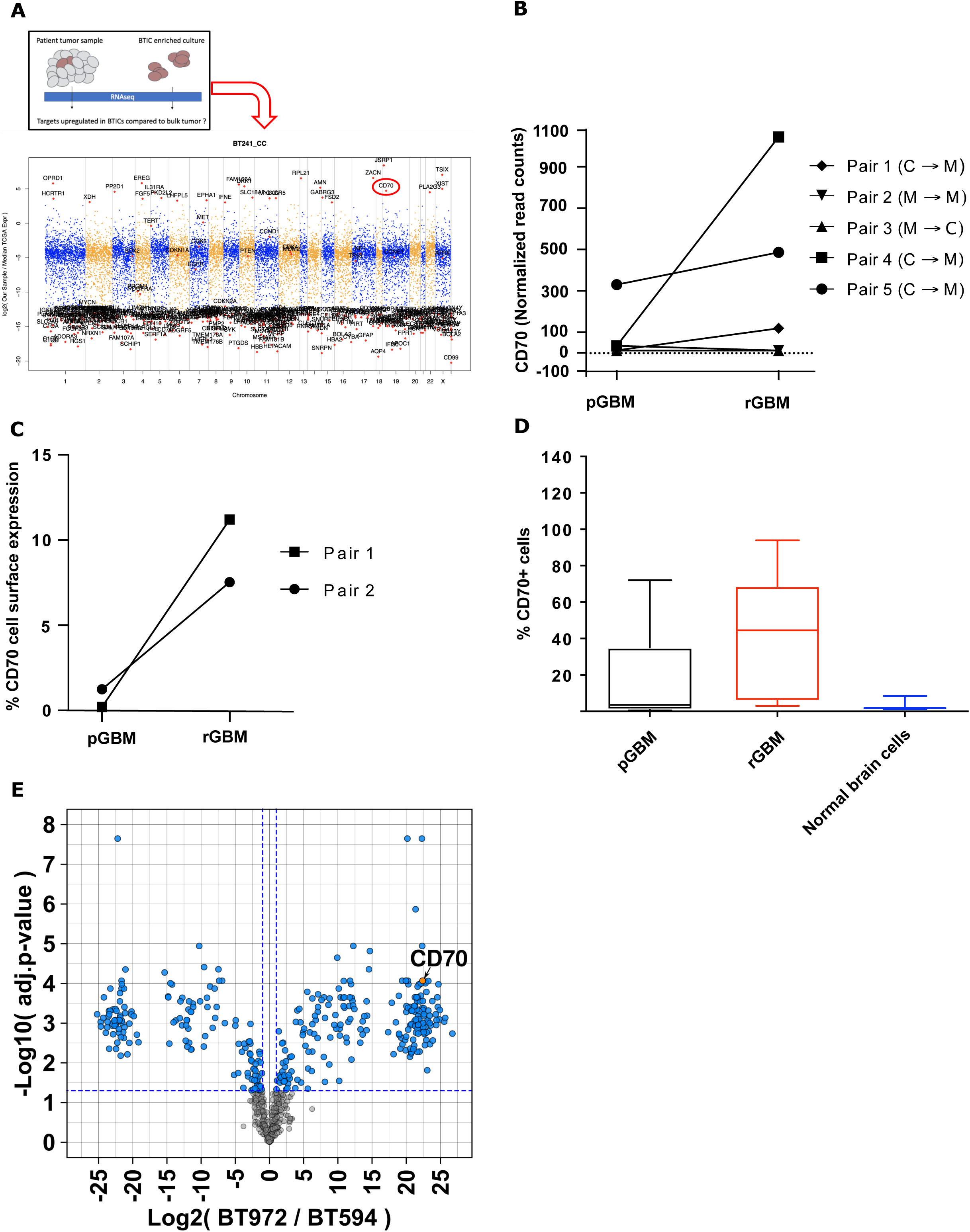
CD70 expression is a relevant marker of recurrent glioblastoma. **A:** Manhattan plot of the top upregulated RNAseq genes in the recurrent patient-derived GBM brain tumor initiating cells (rGBM BTICs) BT241, compared to the publicly available TCGA database, identified CD70 (circled) as a target candidate upregulated in the BTIC subpopulation, compared to tumor bulk. **B:** Analysis of CD70 mRNA levels in the five TGCA primary/recurrent patient matched GBM pairs depicts an CD70 increase in three pairs upon recurrence, alongside a switch towards the Mesenchymal subtype (GBM subtype classification: C, classical; P, proneural; M, mesenchymal). **C:** Among GBM BTICs from Figure 1C, the two in-house matched p/rGBM BTICs pairs display an increase of CD70 cell surface expression upon tumor recurrence. **D:** Box and whiskers representation of a higher CD70 cell surface expression in rGBM BTICs compared to primary (p-) GBM BTICS and normal brain cells (astrocytes, neural stem cells), assessed by flow cytometry (Singh lab brain tumor database). **E:** Volcano plot of the top up- and down-regulated cell surface proteins of pair 2 from Figure 1D, as assessed by glycocapture proteomics.

### CD70 is a key player in GBM maintenance and tumor formation

Given the upregulation of CD70 in GBM, specifically in rGBM, we sought to explore the role that CD70 expression plays in GBM maintenance and progression. We sorted pGBM and rGBM cells as CD70-positive or -negative using FACS analysis and carried out a PrestoBlue proliferation assay. CD70-positive cells demonstrated a significantly increased proliferation capacity compared to their CD70-negative counterparts (Fig2A, SupplFig2A). We next aimed to assess the role of CD70 in sphere formation, a stem-like trait that is typical of BTICs and correlates with self-renewal capacity *in vitro* and tumorigenesis *in vivo* (Hirschhaeuser et al. 2010, Singh et al. 2004). In both CD70^HIGH^ BTIC cell lines, silencing of CD70 using an shRNA knockdown vector led to a significant decrease in sphere formation capacity compared to controls (Fig2B). Given the correlation of sphere formation with tumorigenesis *in vivo*, we investigated whether CD70 silencing limits GBM tumor formation in our patient-derived orthotopic xenograft animal model. We generated CD70 knockdown (shCD70) and control lines (shGFP) of three GBM BTIC lines that naturally express high levels of CD70, and intracranially injected these into immunodeficient mice, as previously described (Singh et al. 2004). We observed a significant decrease in the size of tumors formed by shCD70 cells compared to shGFP controls, as determined by H&E staining (Fig2C-D, SupplFig2C) and MRI imaging (Fig2G). This was further reflected in a significant survival advantage for mice engrafted with shCD70 cells compared to controls (Fig2E and F, SupplFig2D). These findings demonstrate that CD70 plays a key role in recurrent GBM proliferation, tumor formation and survival both *in vitro* and *in vivo*.

**Figure 2:**
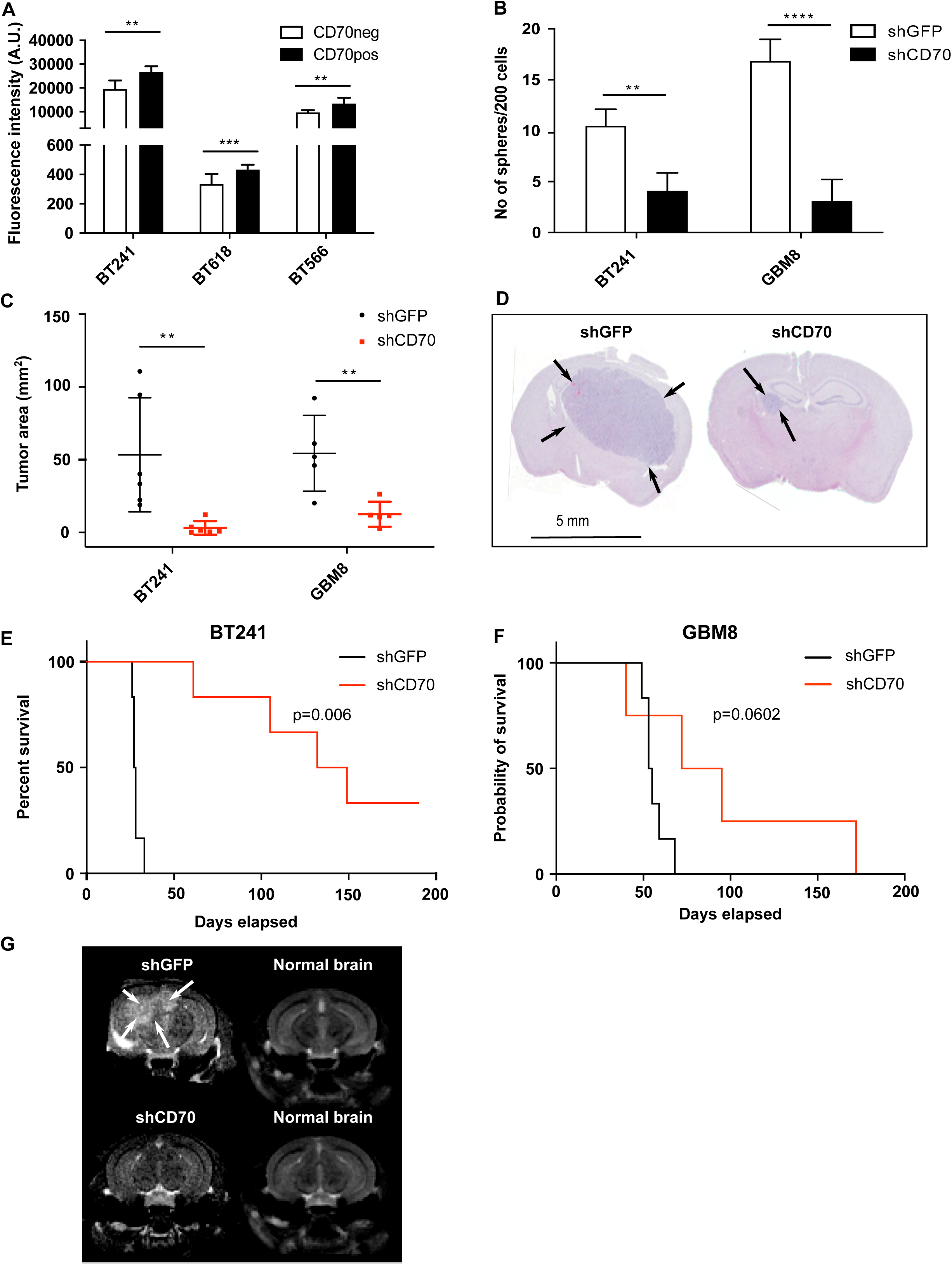
CD70 is a dedicated player in GBM maintenance and tumor formation. **A:** GBM BTICs were sorted into positive and negative populations and proliferation was assessed by PrestoBlue assay **B:** Silencing of CD70 expression by shRNA (shCD70) knockdown and sphere formation ability was assessed compared to shGFP (Control shRNA). **C - F:** Immunocompromised mice (NSG, a minimum of six mice per condition) were intracranially injected with shGFP or shCD70 BTICs. **C, D:** Tumor area of CD70-silenced BTICs compared to control knockdown BTICs was measured using formalin-fixed, H&E-stained mouse brain slices (right, representative picture). **E,F:** Kaplan-Meier survival curves comparing mice engrafted with shCD70 BTICs compared to shGFP BTICs. The two remaining BT241 shCD70 mice at the end of experiment showed an absence of tumor by H&E staining at experimental endpoint (data not shown). **G:** MRI images representative of xenografts from shGFP and shCD70 transduced GBM BTIC line BT241. Right panels are control images of normal mouse brain. (* = p<0.05; ** = p<0.01; *** = p<0.001; **** = p<0.0001)

### CD70 plays a crucial role in cellular programs implicated in tumorigenesis

Previous studies have emphasized the function of CD70 in GBM as it contributes to T-cell apoptosis, and mediates tumor cell migration and invasion, a feature characteristic of mesenchymal-like cells (Ge et al. 2017; Diegmann et al. 2006; Inaguma et al. 2020). To further investigate the role of the CD70 signaling network in GBM, we investigated transcriptional changes and their predicted networks after CD70 silencing using RNA sequencing and subsequent gene set enrichment analysis (GSEA). Using three GBM lines transduced with shCD70 or shGFP, we observed strong downregulation of FOSL1, a gene recently discovered to play a pivotal role in stemness, migration, and EMT (SupplFig3A) (Fiscon, Conte, and Paci, 2018; Feldker et al. 2020). Other slightly downregulated genes included CDH2 (Cadherin 2), PLAUR, and CXCR4; genes known to be associated with the Mesenchymal subtype in GBM and a worse overall prognosis (Gilder et al. 2018; Tao et al. 2020; Yi et al. 2018). OLIG2, a transcription factor commonly associated with the proneural subtype and tumor recurrence (Bouchart et al. 2020, Lu et al. 2016), showed upregulation following CD70 knockdown, while expression of the pro-angiogenic factor VEGFA was depressed, indicating that CD70 may play a role in GBM angiogenesis, a characteristic previously documented in other pathologies, but not in cancer (Simons et al. 2018, Winkels et al. 2017).

GSEA was performed using Gene Ontology (Merico et al. 2010) and MSigDB C2 and C6 gene sets (Subramanian et al. 2005; Liberzon et al. 2011), to gain a deeper understanding of the cellular programs associated with CD70 expression (SupplFig3A). Top modulated pathways showed that silencing CD70 results in downregulation of epithelial-to-mesenchymal transition (EMT) and hypoxia signatures, and upregulation of Interferon type I/interleukin-1 pro-inflammatory signatures (Guarda et al. 2011) (SupplFig3A, B). Hypoxia and EMT pose major hurdles in GBM, as they promote migration of tumor cells further into the brain tissue, while pro-inflammatory signals are often depressed in GBM (Monteiro et al. 2017; Carro et al. 2010; Singh, A, et al. 2010). While these data are limited, they do further implicate the role of CD70 in various processes linked to invasiveness, immunosuppression, and poor prognosis in GBM, as well as angiogenesis and stem-like characteristics of GBM BTICs.

### Generation and characterization of CD70-directed CAR T cells

Adoptive cell therapies have shown great promise in overcoming therapy resistance and providing a more specific targeted therapy in multiple cancers, including in GBM (Bielamowicz, Khawja, and Ahmed, 2013). However, despite significant global efforts to develop these therapies, they have only been approved for B cell malignancies thus far, and have yet to show efficacy in solid tumors such as GBM (O’Rourke et al. 2017). It is believed that this lack of progress is in part due to the immunosuppressive microenvironment of solid tumors, particularly GBM, as well as antigen escape (Sterner and Sterner, 2021).

Given our data implicating CD70 as a key factor in GBM functionality, we tested two distinct in-house fragments antigen-binding (Fabs) for their ability to bind cell-surface CD70, and compared these to commercially available CD70 antibody (Fig4A). The Fab he-Im was used to develop a non-covalently conjugated therapeutic antibody-drug conjugate (ADC) (Fig4B), which we postulated would be advantageous due to the rapid internalization of CD70 upon ligand binding (Adam et al. 2006; McDonagh et al. 2008) (SupplFig4A). We incubated CD70^HIGH^ GBM cells for 72 hours with our ADC, and observed a dramatic cytotoxic effect; an effect not seen in CD70^LOW^ GBM cells or HEK293 control cells, indicating that our ADC is both specific and cytotoxic, and is suitable for developing adoptive cell therapies. Thus, we cloned the scFv region of he-Im into a second-generation CAR linked to a truncated c-Myc tag (Fig4C), and achieved moderate CAR cell-surface expression nine days post-transduction in human T-cells (Fig4D). To determine the efficacy of these anti-CD70 CAR-T (CD70CAR-T) cells, we co-cultured them with CD70^HIGH^ GBM cells. CD70CAR-T cells co-cultured with CD70^HIGH^ GBM cells released significantly more IFN-γ and TNF-α into culture supernatants compared to a control CAR-T (ConCAR-T) (Fig4E, SupplFig 4B, C). Additionally, CD70CAR-Ts demonstrated significant cytotoxicity against numerous CD70^HIGH^ GBM cells at effector to target ratios as low as 1:1 (Fig4F, SupplFig4D). Together, these data indicate that CD70CAR-T cells are capable of mounting a robust and specific immune response against CD70-expressing GBMs.

**Figure 3:**
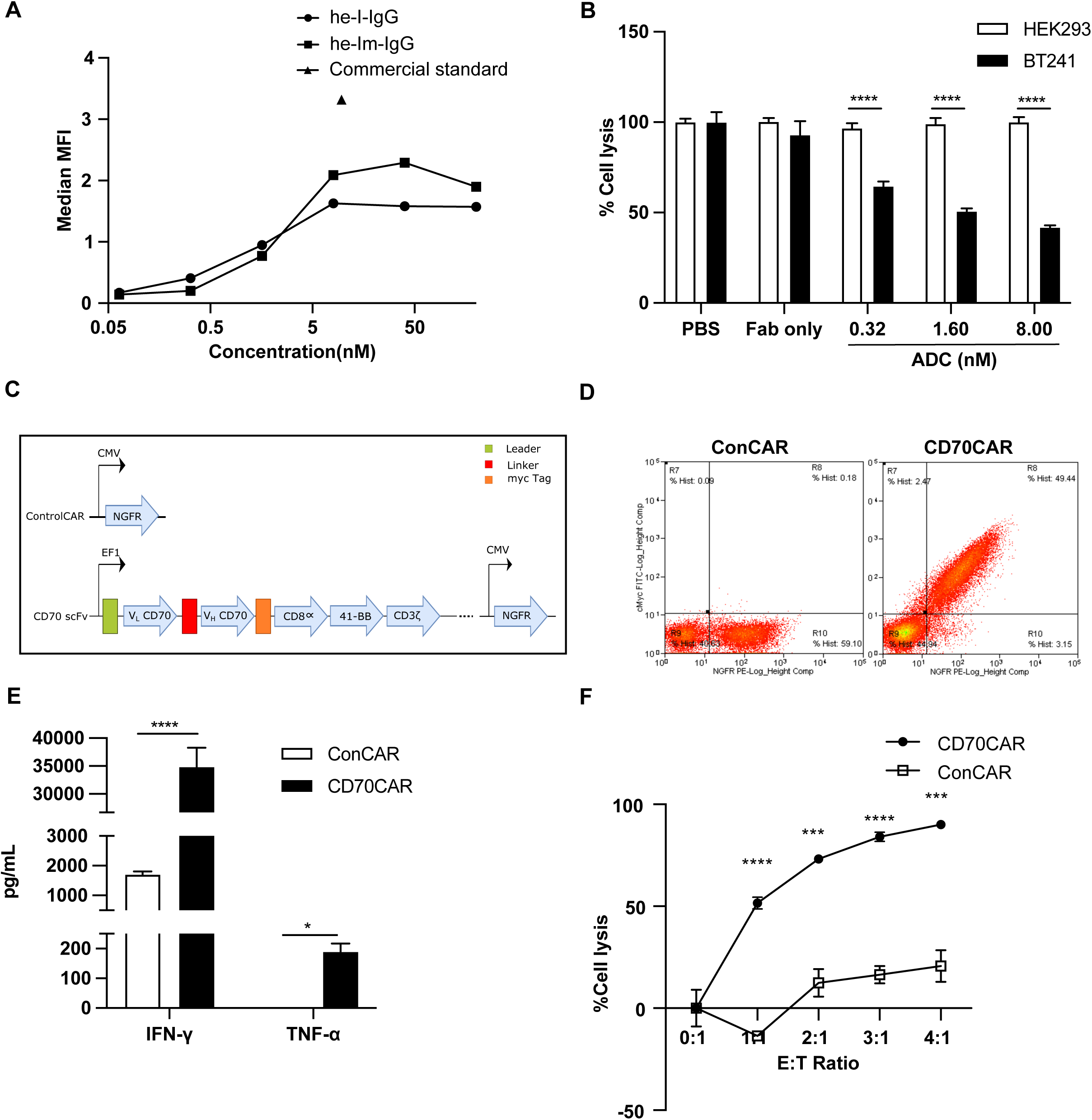
Generation and *in vitro* Characterization of CD70-Specific CAR-T Cells. **A:** Binding curve comparing CD70-specific Fabs to commercial standard antibody. **B:** Anti-CD70 Fab’2 is specific against CD70, assessed by cytotoxicity assay under combination treatment with 2^◦^ADC, against GBM cells expressing high (GBM BTIC BT241) or no (HEK293) CD70. **C:** Schematic representation of CAR structure. **D:** Successful transduction of CAR-T vectors as observed by NGFR+ cells in ConCAR-T cells and NGFR+Myc+ cells in CD70 CAR-T cells, displayed as a representative flow plot. **E:** Testing of CAR-T cell activation; IFN-gamma and TNF-alpha cytokine released during coculture of GBM BTIC BT241 with CD70 CAR-T, compared to ConCAR-T cells, as analyzed by ELISA assay (n=3). **F:** Cytotoxicity assay to assess CD70CAR killing capacity compared to ConCAR after co-culturing for 24 hours, tested at various effector to target (E:T) ratios (n=3). (* = p<0.05; ** = p<0.01; *** = p<0.001; **** = p<0.0001)

**Figure 4:**
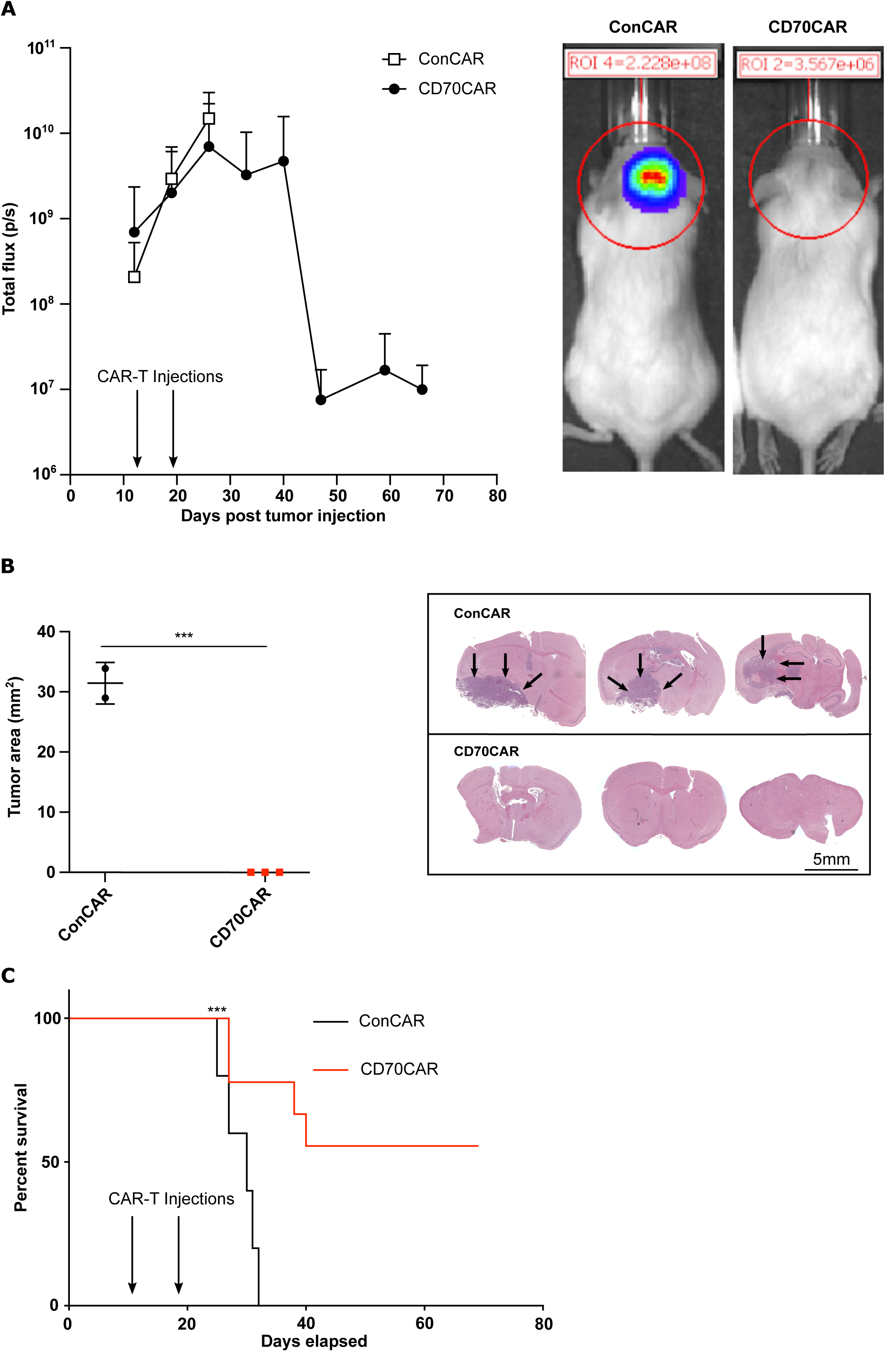
CD70 CAR-T are efficacious against recurrent GBM tumors *in vivo*. NSG mice (at least n=6 per group) were intracranially implanted with 100,000 human BT241 ffLuc GBM cells. Upon successful engraftment, mice were treated with 1×10^6^ CD70CAR-T or ConCAR-T cells, delivered intracranially once a week for two weeks. **A:** CD70 CAR-T treated mice showed decreased tumor signal, as assessed by bioluminescence measurement (right, representative picture of radiance measurement in the region of interest). A lower tumor burden was observed in the CD70 CAR-T group compared to the control group, as measured on B: formalin-fixed, H&E stained mouse brain slices (representative picture on the right), and C: an extended survival (Kaplan-Meier curve) (* = p<0.05; ** = p<0.01; *** = p<0.001; **** = p<0.0001)

Lastly, we assessed the antitumor potential of our CD70CAR-T cells in orthotopically xenografted NODSCID mice, using CD70^HIGH^ BT241 rGBM BTICs. After confirming tumor engraftment using the In Vivo Imaging System (IVIS), we intracranially injected 1M CD70CAR-T or ConCAR-T cells weekly over two weeks. Mice treated with CD70CAR-T cells displayed significantly lower tumor burden, as observed by bioluminescence signal, and a significant decrease in tumor volume as shown by H&E staining, confirming that CD70CAR-T cell-treated mice experience far less tumor growth compared to controls (Fig4A and B). Unsurprisingly based on our previous data, CD70CAR-T cell-treated animals had a significant survival advantage compared to control mice (Fig4C, left panel). Of note is that fact that the majority of animals (5 out of 9) did not display any tumor-related symptoms post-treatment, nor did H&E staining display any presence of tumor at the end of study (Fig4B, right panel). To further validate our CD70CAR-T cell therapy, we reproduced this with another GBM cell line, and observed similar results, indicating that this approach is efficacious in multiple CD70^HIGH^ cell lines (SupplFig5A-C).

### CD70 and its role in the GBM tumor immune microenvironment

CD70 is the only known ligand for the receptor CD27, a TNF receptor superfamily member, and is known to trigger T cell apoptosis and induce exhaustion (Chahlavi et al. 2005), as well as recruit tumor associated macrophages (TAMs) to the GBM microenvironment, contributing to the immunosuppressive nature of GBM (Ge et al. 2017). However, as far as its role in immune system functionality, no evidence has been found to date indicating it is essential (Shaffer et al. 2011). Thus, we elected to investigate the interaction between CD70-expressing GBM cells and CD27-expressing T cells, to see whether there would be any observed effect on T cell viability (Wajant et al. 2016). Additionally, CD70 cleaved from the cell surface and present in the supernatant may act similarly to cell-surface CD70 (Rowley and Al-Shamkhani, 2004). We co-cultured CD70^HIGH^ BT241 cells with CD27+ T cells, and observed a decrease in CD27+ T cell populations, an effect that was not seen when co-culturing with CD70 knockdown BT241 cells, indicating that CD70/CD27 interaction between T cells and GBM cells may initiate apoptotic programs in T cells, adding to the immunosuppressive capacity of GBM, as previously observed (Q. J. Wang et al. 2012) (SupplFig6B). We then cultured CD27+ T cells with supernatant from CD70^HIGH^ GBM cells to observe whether soluble CD70 cleaved into the supernatant could bind CD27+ T cells and initiate downstream apoptotic effects, however, we observed no decrease in CD27+ T cell populations (SupplFig6B).

As seen in the literature (Lens et al. 1997), we observed increased expression of CD70 on activated T cells (Fig5C) and a subsequent decrease in CD70CAR-T cells count and viability, compared to ConCAR-T cells (data not shown). This insinuated that we were observing fratricide between CAR-T cells, as seen previously with other targets (Sánchez-Martínez et al. 2019). Thus, we elected to create a CD70 knockdown CD70CAR-T cell to overcome this problem. For this preliminary model we utilized Jurkat T cells due to their robustness compared to donor-derived T cells, and sorted them by flow cytometry to obtain a CD70^HIGH^ Jurkat cell population. We also created an shCD70 knockdown Jurkat cell population. Both CD70^HIGH^ and shCD70 Jurkats were transduced with CD70CAR construct or ConCAR construct, after which we examined any change in viability and presence of the activation marker CD69 (Fig5E). We were able to demonstrate that CD70^HIGH^ CD70CAR Jurkat cells had decreased viability and increased expression of the activation marker CD69, compared to both ConCAR and shCD70 CD70CAR Jurkats. This indicates that silencing of CD70 may be a viable option for overcoming CD70CAR T fratricide, improving viability of CD70CAR-T cells while having no effect on other T cell functions, as previously shown (Munitic et al. 2013, Kumar et al. 2020).

**Figure 5:**
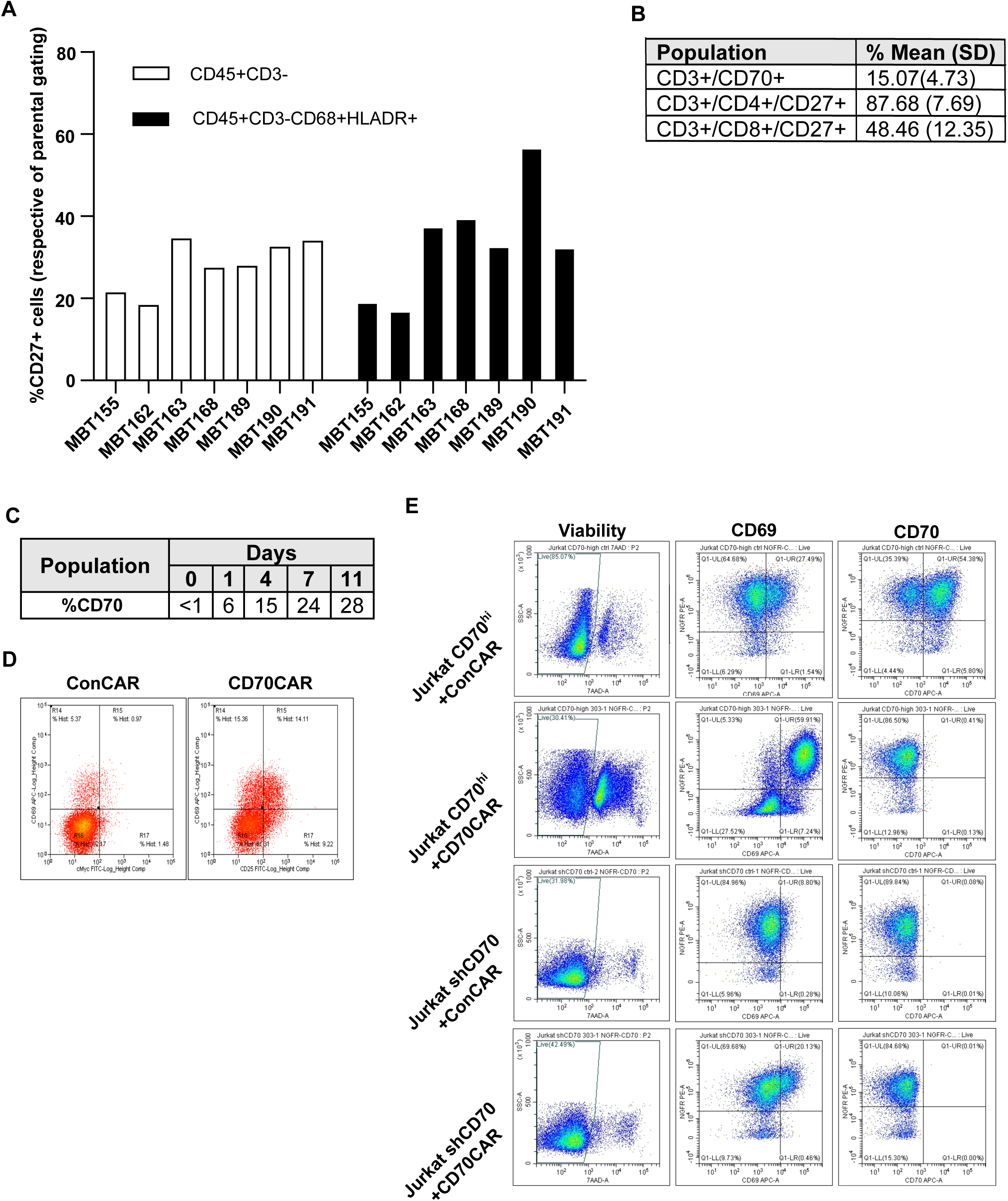
Modelling of CD70 influence on GBM TIME. Tumor immune microenvironment (TIME) cells extracted from patient tumor samples were analyzed by flow cytometry, evaluating the pattern of expression of CD27 in non-lymphoid (CD45+CD3-) and M1 populations (CD45+CD3-CD68+HLADR+). **A, B:** Average expression of CD27 on CD4/CD8 lymphoid population, and CD70 expression on the lymphoid population (CD3+). **C:** CD70 expression kinetics on in-house, activated T cells and **D**: levels of CD69 and cMyc displayed by CD70CAR or ConCAR-T cells nine days post-transduction, evaluated by flow cytometry. **E**: CD70-enriched or -silenced Jurkat cells were transduced with either ConCAR or CD70CAR. After 8 days CD69 and CD70 levels were assessed by flow cytometry. (* = p<0.05; ** = p<0.01; *** = p<0.001; **** = p<0.0001)

According to the literature, cancer cells may utilize the CD70/CD27 interaction to their advantage to modulate both T cells and macrophages to create more immunosuppressive or immune-evasive environments (Ge et al. 2017; Inaguma et al. 2020). Due to the absence of an immune system in our immunodeficient mouse models, we elected to interrogate the GBM tumor immune microenvironment (TIME) using flow cytometry on CD45+ cells extracted from fresh patient-derived tumor samples. From this data we saw that CD27 had a similar expression pattern among non-T-cell populations (CD3-cells) from different tumor samples (Fig6A). Further, CD27 expression was found on putative ‘M1 proinflammatory’ (CD45+CD3-CD68+HLADR+) macrophages to a similar extent (Ma et al. 2010). It is interesting to note that CD27 expression was increased in this particular ‘M1’ population when their corresponding GBM cells were CD70^HIGH^ (i.e., MBT190); a phenomenon not seen when correspond GBM cells were CD70^LOW^ (i.e., MBT162). Interestingly, MBT190 had far fewer CD8+ cytotoxic tumor infiltrating lymphocytes than most samples, including MBT162 (SupplFig6A). In the CD3+ lymphoid population, we found that while CD27 was expressed on the majority of these cells, very few of them expressed CD70 (Fig5A, SupplFig6A), in agreement with existing literature (Sánchez-Martínez et al. 2019). Lastly, we noted that CD27 was highly expressed on CD4 helper T cells, potentially making them more sensitive to CD70^HIGH^ GBMs. Together, these data indicate that multiple CD27/CD70-axis interactions are occurring within the GBM TIME, likely contributing to the low immunogenicity of rGBM.

## DISCUSSION

Since the creation of the Stupp protocol in 2005, few advances have been made in bringing new therapeutics to market for GBM. In addition, resistance to SoC treatment has begun to direct therapeutic investigation towards a small reservoir of cells termed BTICs (Osuka and Van Meir, 2017). BTICs display enhanced self-renewal properties and are believed to be capable of *de novo* tumor formation, and driving tumor recurrence (Osuka and Van Meir, 2017). It is believed that SoC therapy helps drive recurrent tumors by creating a bottleneck, with cells that escape initial chemoradiotherapy driving formation of a therapy-resistant recurrent tumor. Thus, recent endeavors have sought to identify actionable targets on this subpopulation of cells. Numerous clinical trials against these targets have assessed the viability of CAR-T cells against different antigens, dendritic cell vaccines and immune checkpoint inhibitors, amongst others. However, despite many efforts little progress has been made, and most recurrent GBM patients are destined for clinical trials or palliative care.

Here, we utilized a multi-omics approach using our in-house collection of low-passage, patient-derived BTICs, and existing public datasets to identify a GBM cell surface marker of treatment-resistant BTICs. Using this approach, we identified and validated CD70 as a promising therapeutic target in recurrent GBM.

From the literature, CD70 seems to have a confounding role in cancer, both stimulating and suppressing immune response in various cancers (Rowley and Al-Shamkhani, 2004; Aulwurm et al. 2006). With this in mind, we sought not only to validate CD70 as a therapeutic target in GBM, but also to better understand its role in GBM cell maintenance and progression, as well as the tumor immune microenvironment. We demonstrate that CD70 is vital in sphere formation and proliferation of GBM cells, two essential BTIC characteristics which correlate with ability to recapitulate GBM tumors *in vivo*. To follow, we show that CD70 silencing significantly reduce tumor burden and volume, and significantly increases survival time, with some animals showing no signs of disease up until the end of the experiment. These data represent similar findings to that seen in the literature, indicating that CD70 may play a crucial role in tumor-initiating cells in multiple cancers (Liu et al. 2018; Nakamura et al. 2021). To better understand the cellular pathways and programs involving CD70, we carried out GSEA using RNAseq of our CD70-silenced cell lines. We were able to contribute to the functional understanding of CD70 by showing that, as previously noted in RCC and other cancers, CD70 plays a role in controlling hypoxia (Ruf, Moch and Schrami, 2016; Kitajima et al. 2018). However, this is the first time to our knowledge that the regulatory role of CD70 in hypoxia has been demonstrated in GBM specifically, or brain cancer in general. Hypoxia is the consequence of poor vascularization, and thus poor blood delivery, within the tumor. This often promotes cancer cell spreading via invasion so that cells may escape the low-oxygen environment, thereby rendering the tumor diffuse and far more aggressive, a characteristic often seen in recurrent GBM. Based on our GSEA, it is possible that the vascularization factor VEGF is dependent on CD70 expression and the CD70/CD27 signalling axis, leading to, and tumor neoangiogenesis, a role previously established in non-cancer pathologies (Simons et al. 2018). Our analysis also revealed that CD70 silencing appeared to depress pathways related to EMT signaling, a program that is linked to a more aggressive cancer capable of therapy evasion in multiple cancers, including GBM (Tulchinsky et al. 2019; Zheng et al. 2015; Perotti et al. 2019; Tang et al. 2016).

By defining the tumorigenic significance of CD70 in GBM, we sought to develop potential therapeutic modalities directed against CD70. We developed a CAR against CD70 and demonstrated its efficacy both *in vitro* and *in vivo*, where it displayed high specificity for CD70, and conferred significantly extended survival in our orthotopically xenografted animal models, with some animals experiencing complete remission. CD70CAR-T cell efficacy varied slightly *in vivo*, which may be due to the extensive heterogeneity of GBM, particularly in early passage GBM BTICs which recapitulate intratumoral heterogeneity quite well. Changes in associated clonal dynamics which arise with therapeutic pressure may generate therapy-resistant CD70^LOW^ subpopulations with antigen escape, as has been seen in the clinic (Kim et al. 2015). While antigen escape is a major problem at the moment, particularly as not all BTICs uniformly express CD70 (SupplFig1B) or any other single targetable protein, it is may be possible to overcome this issue using a poly-therapeutic strategy. Examples of such a combinatorial strategy include a bispecific anti-CD70/SIRPα antibody which outperforms individually delivered antibodies in models of human Burkitt’s lymphoma, allowing for co-targeting of both tumor cells and tumor-associated macrophages in the TIME (Ring et al. 2017, https://doi.org/10.1073/pnas.1710877114). We also detected a population of CD27+ pro-inflammatory putative ‘M1’ macrophages in our profiling of patient-derived rGBM TIME samples, indicating that while the TIME is immunosuppressive, it does contain traditionally anti-tumor components. CD27 is a marker of highly immunosuppressive Tregs (Starzer and Berghoff, 2020), and it is possible that tumor associated macrophages and other members of the TIME may exert some of their immunosuppressive effects through CD27/CD70 interactions, inducing a more immunosuppressive phenotype in tumor infiltrating lymphocytes (Jacobs et al. 2015). Among the TCGA matched pairs, we noticed a trend of subtype switching to the mesenchymal subtype upon recurrence, as has been observed in the literature (Segerman et al. 2016, Liu et al. 2018), and increased CD70 expression correlated with the mesenchymal subtype as well. The mesenchymal subtype is associated with a more aggressive tumor, as well as invasion, therapy evasion and a poorer prognosis (Q. Wang et al. 2017; Pich, C et al. 2016), and has been shown to harbour a stronger immunosuppressive microenvironment. While initially this may seem to be negative as far as the impact cell therapies might have, some studies suggest the opposite, and that immunotherapies may have a more drastic impact on these tumors (Haddad et al. 2020; Chen and Hambardzumyan, 2018).

In accordance with the literature, after silencing of CD70 expression in various GBM cell lines, we found that CD70 repression results in downregulation of IFN-α and IL-1, both of which play an important role in the Th1 lymphocyte response, and are known for their pro-inflammatory role in tumors (Biron, 1998; B. S. Kim et al. 2012). In conjunction with other work highlighting CAR-T cells’ ability to induce inflammation (Gajewski et al. 2017), we see the potential for CD70CAR-T cells to convert the GBM microenvironment from immunologically “cold” to “hot”, eliciting an anti-tumor immune response of endogenous effector cells. It has previously been shown that certain aspects of T cell functionality are dependent on IFN-α and CD70 (Allam et al. 2014), although to our knowledge this is the first time that this dependence has been shown in cancer cells.

Our work, as well as recent work from others (Ge et al. 2017), maintain that CD70 is a promising target for rGBM, and a better understanding of the role of CD70 in the GBM TIME is needed. In particular, we advocate for determining the role that CD70 plays in initiating GBM recurrence and potential mechanisms of therapy evasion. While the literature shows conflicting results regarding whether CD70 plays a pro- or anti-tumorigenic effect in cancer, we postulate a novel mechanism, through which continued stimulation of the CD70/CD27 axis leads to continuous T cell activation by GBM cells, and subsequent exhaustion within the T-cell compartment, as previously observed in models of HIV (Tesselaar et al. 2003).

As has been investigated in the literature, CD70 is present on the cell surface of T-cells, however, it does not play a functional role, and as a result, other groups have noted fratricide in their CD70 CAR-T cell populations (Wang, Q. J. et al. 2018). Based on these reports, as well as our own observations collected from our CD70 CAR-T cell populations, we developed a model of CD70-CAR T cell fratricide using Jurkat cells, which may be used as a platform for future studies in adoptive cell therapy. Reasons for the observed decrease in viability of Jurkat cells, though not assessed experimentally here, have previously been speculated upon by colleagues using CAR-transduced Jurkat cells (Raikar et al. 2017), reporting that cell death occurs due to prolonged activation-induced paracrine and autocrine interactions. Compared to the flow cytometry-sorted CD70-enriched Jurkat cells used in our experiments, CD70 expression is relatively scarce on activated T cells, indicating that fratricide would occur to a far lesser extent on natural PBMC-derived CD70 CAR-T cells. In our hands, only small variations in cell viability were observed in our CD70 CAR-T cell experiments. Nonetheless, others recently reported an additional fitness benefit in a CD70 knockout CD70 CAR-T cell, which conferred various functional benefits including increased proliferation and cytotoxicity, and was far more advantageous than other obvious knockout targets such as PD-1 and LAG3 (Dequeant et al, 2021). Thus, we highly encourage future studies exploring any potential benefit a CD70KO CD70 CAR-T cell may display in GBM, where the TIME is notoriously difficult to overcome. Targeted delivery of the CAR construct directly into the CD70 locus would disrupt CD70 expression, preventing undesirable stimulation and enhancing proliferation and cytotoxicity of CD70 CAR-T cells (Kumar et al. 2020). Strategies involving targeted gene delivery have already been applied to an array of other targets to optimize adoptive cell therapies (Cooper et al. 2018).

Here, we investigated the functional benefit that CD70 expression confers in GBM cells, and the implied influence that CD70 expression may have on interactions with the immunosuppressive landscape. We employed a reverse translational approach (Goswami et al. 2020) to determine CD27’s expression pattern – CD70s only known receptor – in different compartments of the GBM TIME. This highlights the influence that CD70-expressing GBMs may have on their microenvironment by leveraging this interaction, and how utilizing a ‘double jeopardy’ therapeutic strategy – targeting both GBM cells and immunosuppressive microenvironment cells – may result in highly potent anti-tumor activity. Considering our data and that of recent clinical trials targeting CD70 by systemic administration (Riether et al. 2020), we believe intracranially delivered CD70 CAR-T therapy holds great promise, and should be explored alone and in conjunction with TIME-targeting therapeutics.

## ACKNOWLEDGMENTS

M.S. was supported by MITACS fellowship along with Longbow therapeutics. S.K.S. is supported by the Terry Fox Program Project grant and McMaster University Department of Surgery. We thank Dr. Mary Haak-Frendscho and Dr.Alan Wahl for providing us with CD70 antibodies.

## Supplementary Figures

**Supp Figure 1: CD70 expression on in-house primary and recurrent GBM samples.**

**A.** Manhattan plot of the top upregulated RNAseq genes in patient-derived GBM brain tumor initiating cells (GBM BTICs) BT698, BT956, BT618, compared to the publicly available TCGA database, identified CD70 (circled in red) as a target candidate upregulated in the BTIC subpopulation, compared to tumor bulk. **B:** CD70 cell surface protein expression, assessed by flow cytometry cell surface staining on multiple in-house primary and recurrent cell lines, as well as normal human cell lines.

**Supp Figure 2: Knockdown of CD70 improves survival *in vivo*.**

**A:** Proliferation of sorted CD70-positive and -negative cells from BT698, BT428, BT458 GBM cultures was assessed using a PrestoBlue assay. **B:** Cell surface CD70 expression after shRNA knockdown in three CD70^HIGH^ GBM lines, as assessed by flow cytometry. **C:** Tumor area from animal-engrafted tumors of shCD70 GBM4 cells compared to control. **D:** *Kaplan-Meier* curve displaying survival of mice engrafted with shCD70 GBM4 cells compared to shGFP GBM4 controls.

**Supplementary Figure 3: CD70 influence on the MES subtype and associated machinery.**

**A, B**: RNA sequencing of CD70 silenced GBM BTIC cells BT241, GBM8 and GBM4 permitted the gene set enrichment analysis (GSEA) (A) and Cytoscape Node map (B) depicting circuitries under CD70 dependence.

**Supp Figure 4: Generation and *in vitro* Characterization of CD70-Specific CAR-T Cells.**

**A:** Internalization studies showing Fab presence on the surface of BT241 cells after a 2 hour incubation. **B, C:** IFN-gamma and TNF-alpha release during coculture of CD70 CAR-T or ConCAR with GBM BTICs lines GBM4, GBM8, or BT935 CD70 CAR-T, at effector to target (E:T) ratios of 1:1, as analyzed by ELISA (n=3). **D:** Cytotoxicity assay assessing the killing capacity of CD70CAR-T cells on GBM8 BTICs compared to control, after co-culturing for 24 hours, as tested at various E:T ratios (n=3).

**Supp Figure 5: CD70 CAR-Ts are efficacious against recurrent GBM8 cells *in vivo*.**

NSG mice (at least n=6 per group) were intracranially implanted with 100,000 human GBM8 ffLuc GBM cells. Upon successful engraftment, mice were treated with 1×10^6^ CD70CAR-T or ConCAR-T cells, delivered intracranially once a week for two weeks. **A:** CAR-T treated mice a lower tumor burden in the CD70 CAR-T group compared to Control group as by IVIS imaging. **B:** CD70CAR-T treated mice also showed higher survival rate compared to that of the ConCAR-T cohort. **C:** Tumor burden was assessed in the CD70 CAR-T group compared to Control group, as measured using formalin-fixed, H&E stained mouse brain slices (representative picture on the right).

**Supp Figure 6: Modelling CD70s influence on the GBM TIME.**

**A.** Tumor immune microenvironment (TIME) cells extracted from patient tumor samples were analyzed by flow cytometry, evaluating the pattern of expression of CD27 in non-lymphoid cells (CD45+CD3+) as well as cytotoxic T cell infiltration (CD8+ in CD45+CD3+). **B.** Viability of CD3+ or CD3+CD27+ cells were assessed after co-culture with cells only or supernatant only from i) control knockdown BT241 cells; ii) one of two constructs that produce CD70KD BT241 cells.

---

| **Sample** | **Age** | **Sex** | **Primary/Recurrent (P/R)?** |
| --- | --- | --- | --- |
| BT241 | 66 | F | R |
| BT972 | 53 | M | R |
| BT594 | 50 | M | P |
| BT618 | 67 | F | R |
| BT566 | 55 | F | R |
| BT155 | 66 | F | P |
| BT162 | 57 | F | P |
| BT163 | 56 | M | P |
| BT168 | 46 | F | P |
| BT189 | 67 | F | P |
| MBT190 | 27 | F | R |
| BT191 | 50 | F | P |
| BT956 | 70 | F | P |
| BT698 | 57 | F | P |
| MBT225 | 27 | M | R |
| MBT173 | 69 | F | R |
| BT566 | 55 | F | R |
| MBT103 | 52 | F | P |
| MBT121 | 67 | F | P |
| BT458 | 81 | M | P |
| BT428 | 63 | F | P |
| BT954 | 65 | M |  |
| BT799 | 77 | F | P |
| MBT63 | 68 | F | P |
| MBT27 | 56 | M | P |
| BT935 | 53 | F | P |
| BT459 | 60 | M | P |
| BT778 | 54 | F | P |

| Ensembl ID | Gene symbol | Log (FC) |  |  | Combined Differential Gene Expression |  |
| --- | --- | --- | --- | --- | --- | --- |
|  |  | BT241 | GBM8 | GBM4 | Log (FC) | P-value |
| ENSG00000175592 | FOSL1 | -3.0876903 | -2.9685466 | -1.4827605 | -1.3302582 | 0.002764603 |
| ENSG00000205927 | OLIG2 | 0.99188628 | 1.51882472 | 2.02962725 | 0.95394172 | 0.027641632 |
| ENSG00000112715 | VEGFA | -1.079047 | -2.993489 | -0.2838893 | -0.8847545 | 0.040829495 |
| ENSG00000170558 | CDH2 | -1.8132171 | -0.4377818 | -1.1326002 | -0.633147 | 0.148275349 |
| ENSG00000011422 | PLAUR | -2.0416545 | -1.1341974 | -2.2947466 | -0.4965695 | 0.206493437 |
| ENSG00000121966 | CXCR4 | -1.5615955 | -2.8821719 | -1.3521936 | -0.361023 | 1 |
| ENSG00000026508 | CD44 | -0.6256976 | -0.0583328 | -1.5159146 | -0.3010297 | 0.482572636 |
| ENSG00000181449 | SOX2 | -0.0175167 | -0.8038392 | -0.905513 | -0.3279201 | 0.417025332 |

