## Supplementary Figures and Tables for "Hitting more birds with one stone: CD70 as an actionable immunotherapeutic target in recurrent glioblastoma"

**A**

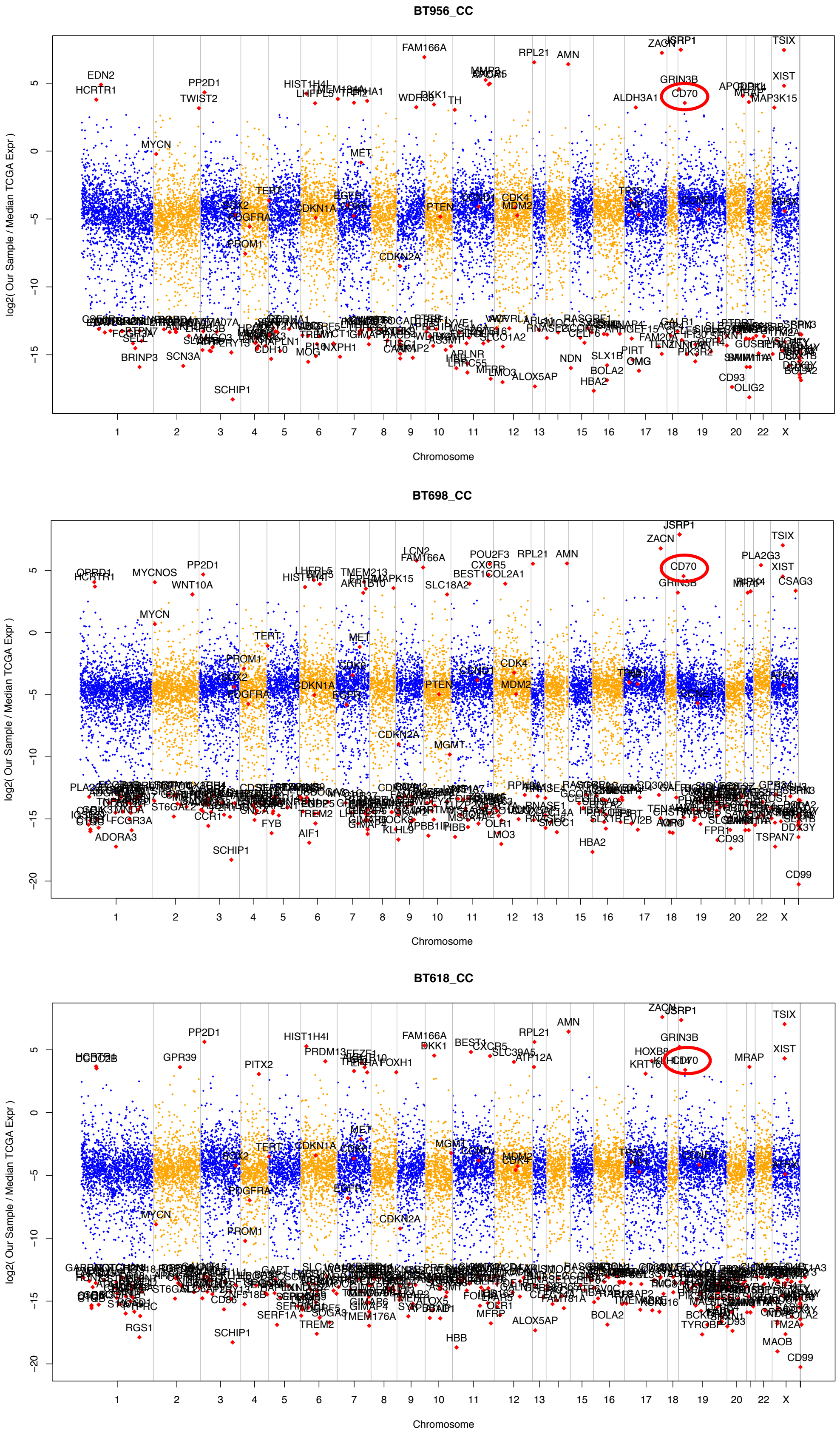


**B**

| Patient derived BTIC lines | %CD70+ cells |
|---|---|
| BT241 | 94 |
| MBT190 | 69 |
| MBT225 | 68 |
| BT618 | 46 |
| MBT173 | 41 |
| BT972 | 11.3 |
| SB2b | 4 |
| BT566 | 3 |
| GBM4 | 72 |
| GBM8 | 70 |
| BT698 | 65 |
| RN1 | 35 |
| MBT103 | 35 |
| MBT121 | 12 |
| BT458 | 9 |
| BT428 | 5 |
| BT954 | 4 |
| WK1 | 3 |
| BT799 | 2.2 |
| GBM123 | 1.5 |
| MBT63 | 1.3 |
| MBT27 | 1 |
| BT935 | <1 |
| BT594 | 0.5 |
| BT459 | <1 |
| BT778 | <1 |
| NSC195 | 1.8 |
| NSC201FT | 8.5 |
| Astrocytes | 1.2 |

**Sup Figure 1**

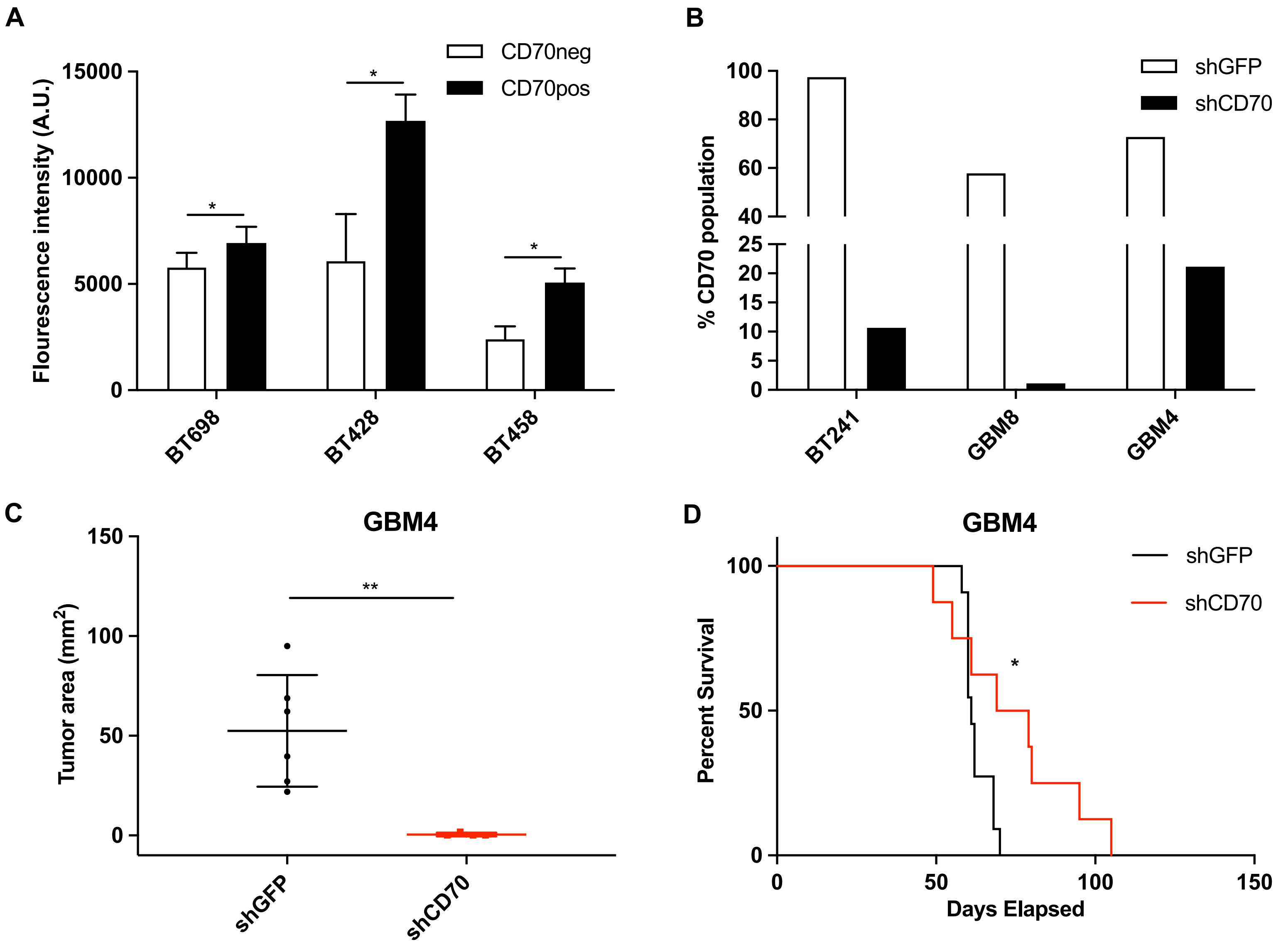
A
CD70neg
CD70pos
Flourescence intensity (A.U.)
15000
10000
5000
0
*
*
*
BT698
BT428
BT458
B
shGFP
shCD70
% CD70 population
100
80
60
40
25
20
15
10
5
0
BT241
GBM8
GBM4
C
GBM4
Tumor area (mm2)
150
100
50
0
**
shGFP
shCD70
D
GBM4
shGFP
shCD70
Percent Survival
100
50
0
*
0
50
100
150
Days Elapsed

Sup Figure 2

**A**

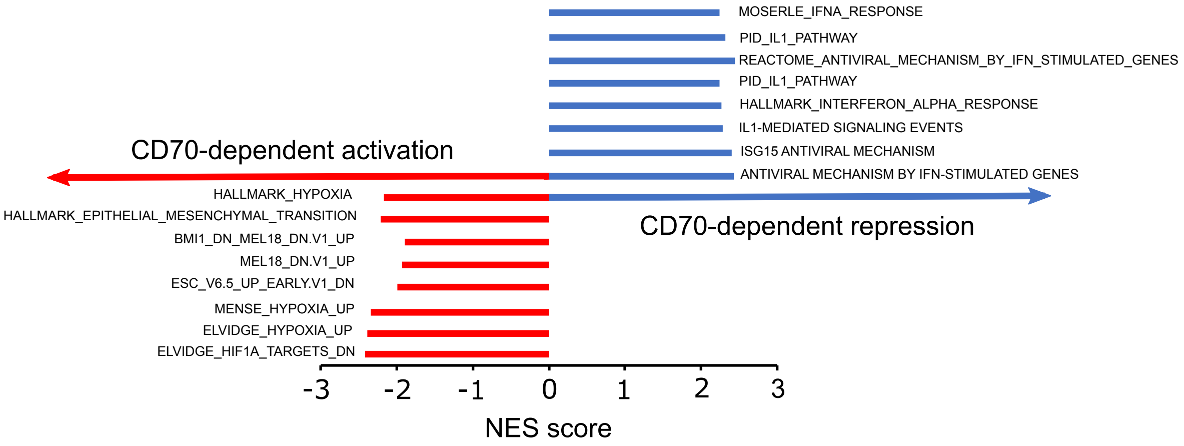


**B**

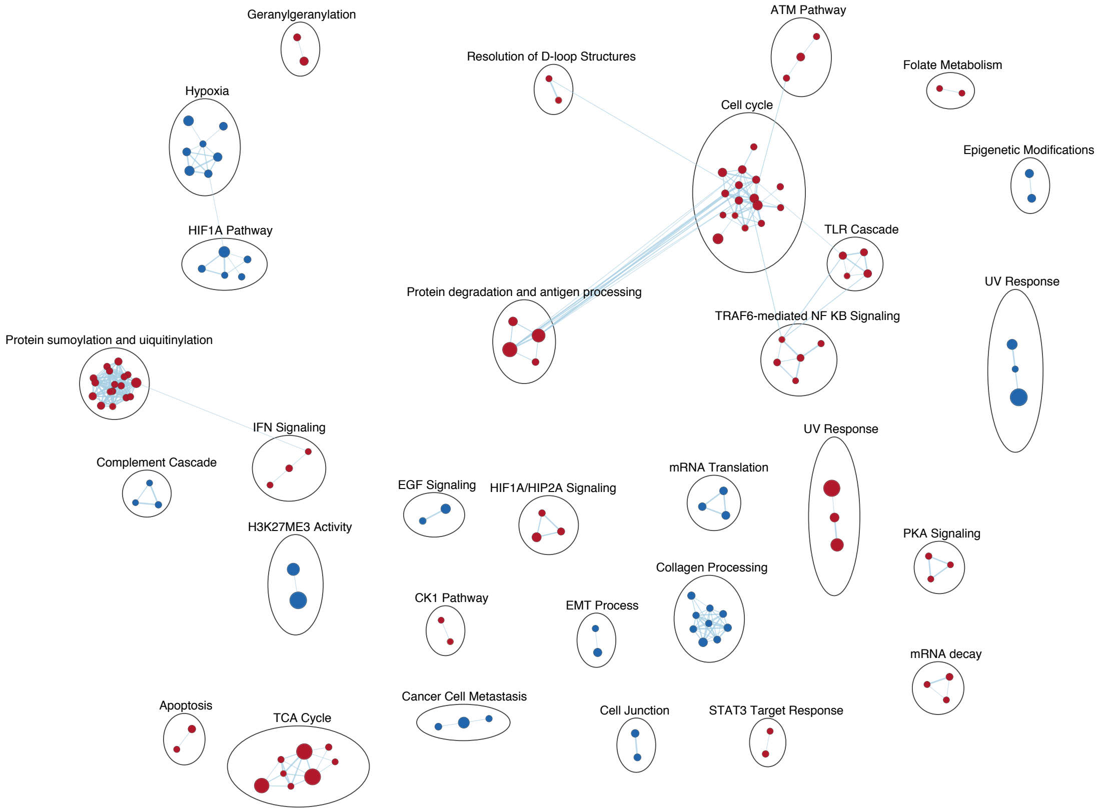


**Sup Figure 3**

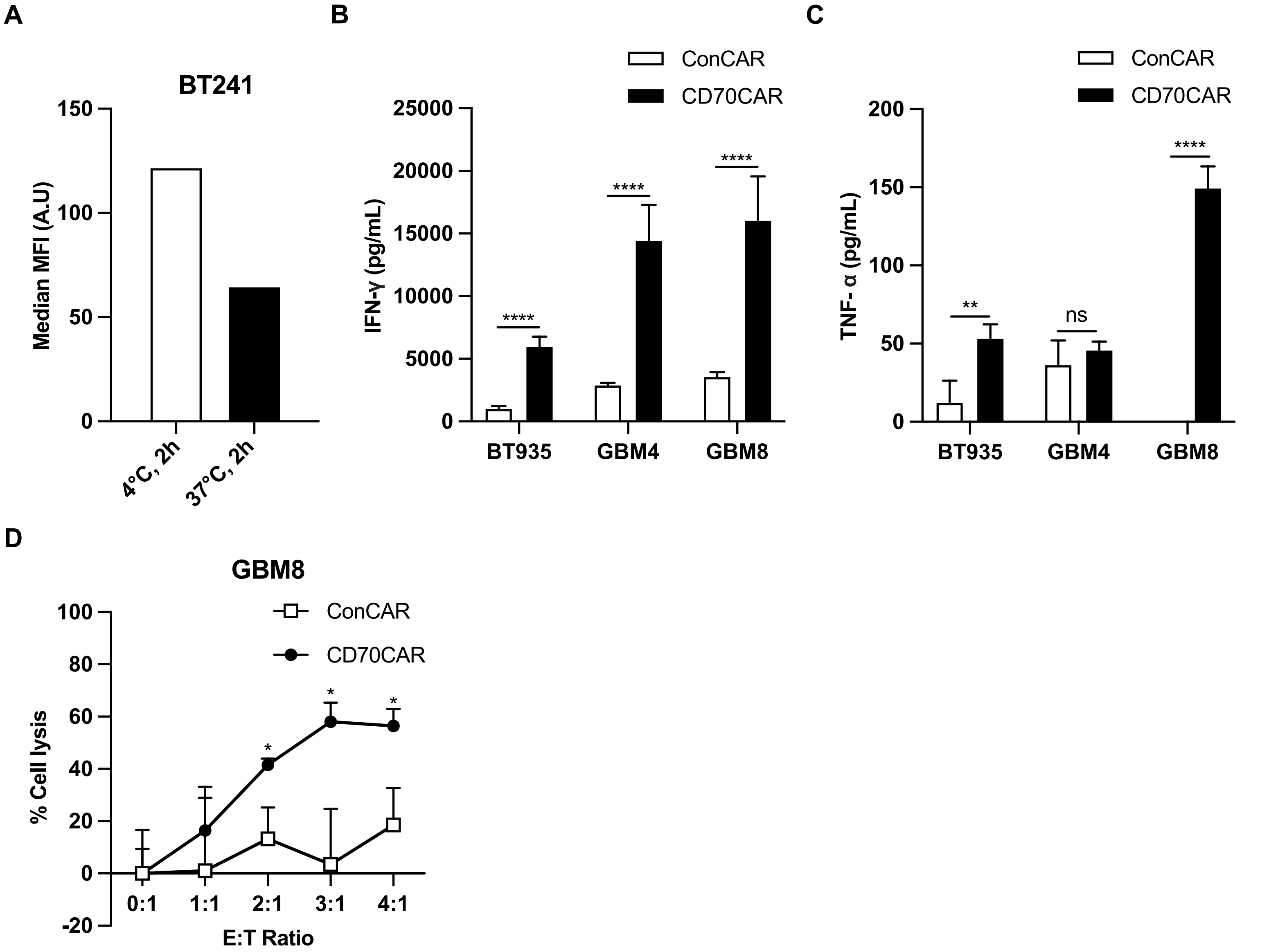
A
BT241
Median MFI (A.U)
150
100
50
0
4°C, 2h
37°C, 2h
B
ConCAR
CD70CAR
IFN-γ (pg/mL)
25000
20000
15000
10000
5000
0
****
****
****
BT935
GBM4
GBM8
C
ConCAR
CD70CAR
TNF- α (pg/mL)
200
150
100
50
0
**
ns
****
BT935
GBM4
GBM8
D
GBM8
ConCAR
CD70CAR
% Cell lysis
100
80
60
40
20
0
-20
*
*
*
0:1
1:1
2:1
3:1
4:1
E:T Ratio

Sup Figure 4

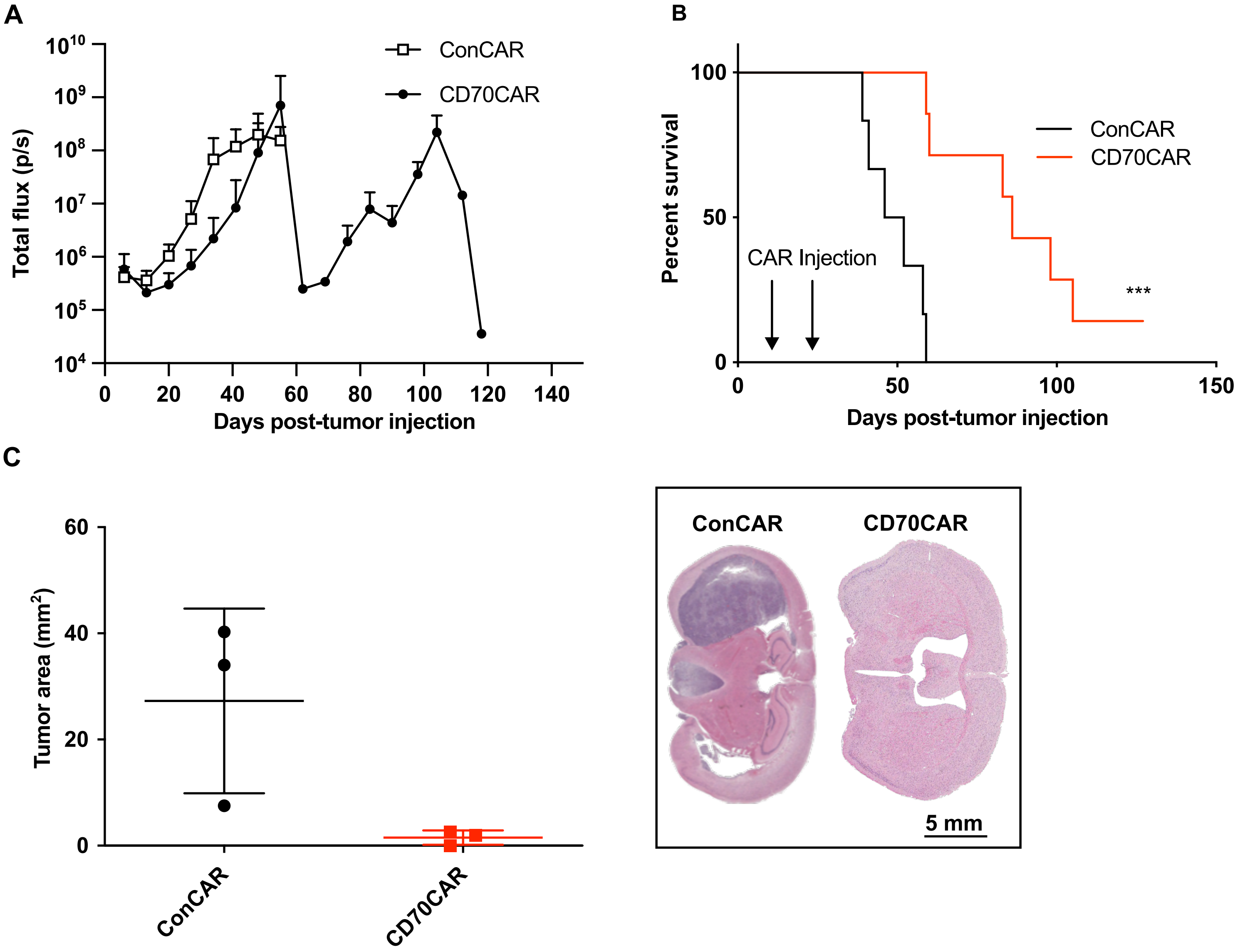
A
$10^{10}$
$10^{9}$
$10^{8}$
$10^{7}$
$10^{6}$
$10^{5}$
$10^{4}$
Total flux (p/s)
ConCAR
CD70CAR
0
20
40
60
80
100
120
140
Days post-tumor injection
B
100
50
0
Percent survival
ConCAR
CD70CAR
CAR Injection
***
0
50
100
150
Days post-tumor injection
C
60
40
20
0
Tumor area ($mm^2$)
ConCAR
CD70CAR
ConCAR
CD70CAR
5 mm

Sup Figure 5

**A**

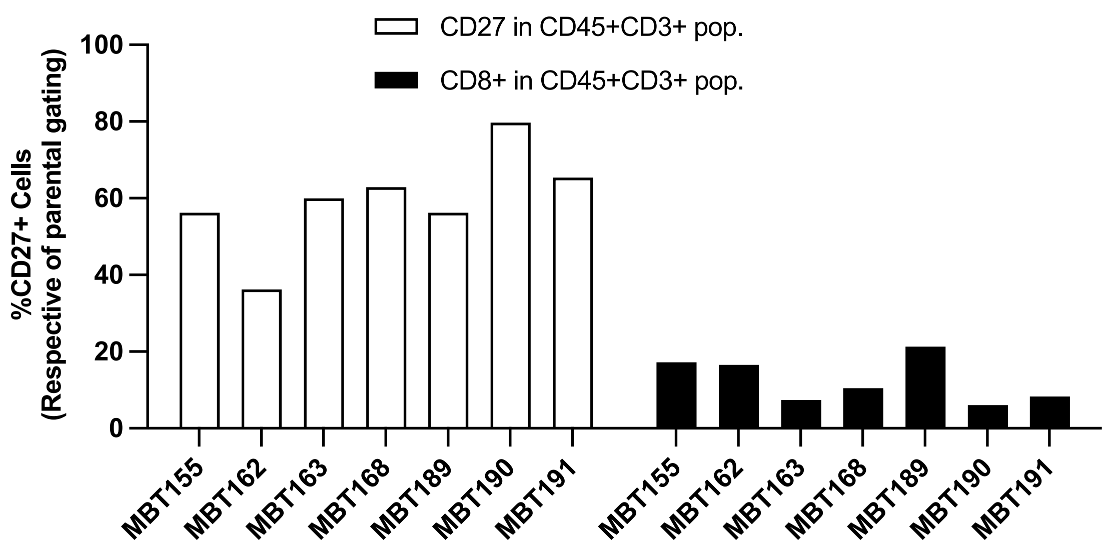


**B**

| Experimental setup | %CD3+ | %CD3+/CD27+ |
|---|---|---|
| T cells alone | 84 | 78 |
| T cells + BT241 crAAVS1 (supernatant) | 76 | 78 |
| T cells + BT241 crCD70-A (supernatant) | 76 | 78 |
| T cells + BT241 crCD70-B (supernatant) | 78 | 78 |
| T cells + BT241 crAAVS1 (cells) | 77 | 43 |
| T cells + BT241 crCD70-A (cells) | 77 | 79 |
| T cells + BT241 crCD70-B (cells) | 80 | 76 |

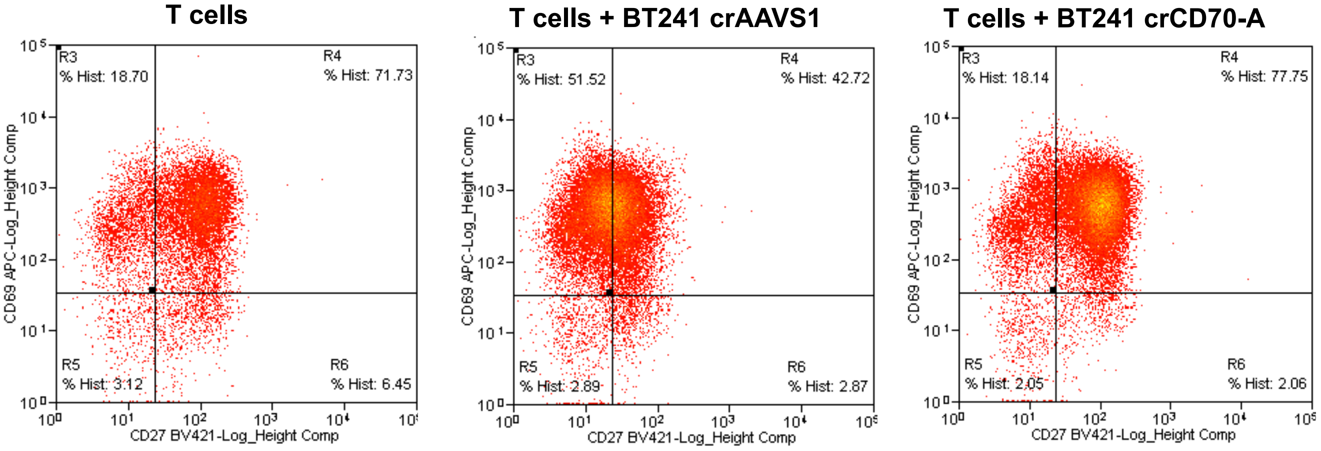


Sup Figure 6
